# Decoding glycosylation potential from protein structure across human glycoproteins with a multi-view recurrent neural network

**DOI:** 10.1101/2024.05.15.594334

**Authors:** Benjamin P. Kellman, Julien Mariethoz, Yujie Zhang, Sigal Shaul, Mia Alteri, Daniel Sandoval, Mia Jeffris, Erick Armingol, Bokan Bao, Frederique Lisacek, Daniel Bojar, Nathan E. Lewis

## Abstract

Glycosylation is described as a non-templated biosynthesis. Yet, the template-free premise is antithetical to the observation that different N-glycans are consistently placed at specific sites. It has been proposed that glycosite-proximal protein structures could constrain glycosylation and explain the observed microheterogeneity. Using site-specific glycosylation data, we trained a hybrid neural network to parse glycosites (recurrent neural network) and match them to feasible N-glycosylation events (graph neural network). From glycosite-flanking sequences, the algorithm predicts most human N-glycosylation events documented in the GlyConnect database and proposed structures corresponding to observed monosaccharide composition of the glycans at these sites. The algorithm also recapitulated glycosylation in Enhanced Aromatic Sequons, SARS-CoV-2 spike, and IgG3 variants, thus demonstrating the ability of the algorithm to predict both glycan structure and abundance. Thus, protein structure constrains glycosylation, and the neural network enables predictive *in silico* glycosylation of uncharacterized or novel protein sequences and genetic variants.

## Introduction

Glycosylation is difficult to study as the one supposedly non-templated biopolymer.^1^ Unlike RNA, DNA, and proteins, glycan sequences are understood to be determined by local metabolic and enzymatic conditions, including the availability of charged nucleotide sugars, enzyme availability, Golgi localization, and substrate competition.^2^ These well-supported claims do not explain how different glycosylation sites within one protein are consistently differentially glycosylated; a phenomenon called “microheterogeneity.”^3^

Indications of protein structure bounded biosynthesis for glycans has existed for decades. After the N-glycosylation sequon (NX[S/T]) was defined, proximal-amino acid variation was found to impact glycosylation complexity,^4–6^ occupancy,^7^ efficiency,^8^ and glycan class.^9^ Conversely, amino acid sequence alignments of similarly glycosylated glycosites suggest the presence of glycosite-flanking sequence conservation.^10^ In influenza and HIV, variation in glycosylation and genetic variation proximal to glycosites can facilitate immune evasion.^11,12^ Examples of how the protein context can constrain glycosylation include observations of higher-order structures such as β-sheets and α-helices,^13^ accessibility,^14–16^ and glycosylation kinetics,^17–20^ all of which impact glycan structure. We quantified associations between glycan substructures and local protein structure, showing that protein structural constraints can predict glycosylation. Together, these protein-glycan relations form a more comprehensive framework we call bounded biosynthesis, wherein glycosylation is bounded by both metabolic conditions and genome-encoded protein structural constraints.^21^ That study describes protein structure as a major determinant of glycosylation, but there is a need to functionalize the proteomic bounds on glycosylation such that it can be leveraged with ease to predict glycosylation from protein structure.

Machine learning can be applied to the complex structures of glycans for the analysis of glycan structure, function, and classification. For example, natural language processing can encode glycans longitudinally from the reducing end.^22,23^ The SweetTalk glycan embedding recapitulated both antigenic glycans and microbial pathogenicity and phylogeny. Another study leveraged the branched nonlinear glycan structure to scaffold graph convolutional neural networks.^24^ SweetNet identified glycan targets of viral lectins. Beyond glycan embedding, biosynthetic constraints and outcomes have been modeled using neural networks.^25^ Previous attempts have been made to relate glycan branching with glycosite-proximal protein structure.^26^ In the absence of meaningful embeddings and biosynthetic-substructure decomposition like SweetNet and GlyCompare,^27^ previous observations were limited to the association between surface accessibility and glycan complexity. With these new embeddings and the knowledge that glycan biosynthesis is a protein structure guided process, we can now functionalize protein-based glycan predictions.

Here we present the Interloping Saccharide Neural Network Extrapolation (InSaNNE) model, which predicts N-glycosylation from glycosite-proximal protein features. Using long short-term memory (LSTM) units,^28^ a type of recurrent neural network, we analyze glycosite-proximal amino acids and leverage the functional and biosynthetic glycan encodings of SweetTalk, SweetNet, and GlyCompare to generate an accurate mapping of glycan structure to protein sequence and structure. We train and validate our glycosite-glycan pairing model on empirically observed site-specific glycosylation. The model is trained using data from UniCarbKB^29^ and validated using more extensively curated data from GlyConnect^30^. We further validate our predictions on important glycosylation events on the coronavirus spike protein, immunoglobulin, and the enhanced aromatic sequon. All N-glycan predictions are integrated in GlyConnect for easy access. With InSaNNE, we leverage the new bounded biosynthesis paradigm to open glycobiology to everyone by predicting expected and differential glycosylation onto their proteins of interest.

## Results

### Graph convolutional neural networks accurately predict glycan-glycosite pairs

We developed a model to predict the presence of specific glycans given the flanking amino acid sequence at N-linked glycosylation sites. Specifically, glycan structures can be ranked to indicate the most feasible glycosylation events at a glycosite of interest. To train, validate, and test the model, we collected and annotated 1,721 unique glycosylation events across 75 human glycoproteins from UniCarbKB^29^ wherein glycan structure was previously fully determined (see Methods). The model incorporates modules that analyzed both glycan structures (**Figure 1**a) and the protein sequences (**Figure 1**b). To analyze the protein sequences, we used long short-term memory (LSTM) units,^28^ a recurrent neural network module effective at modeling protein structure by asserting language-like processivity^31^ (**Figure 1**b). Both sequence-proximal (glycosite-flanking) and spatially proximal (within n-Angstroms) protein features are important for predicting feasible glycosylation. We examined two separate LSTM-based modules into our model for analyzing the sequence-proximal and spatially proximal amino acids, separately. For the analysis of the glycan component, we tested three glycan embeddings: (1) a fully connected neural network using GlyCompare glycan substructure features^27^ as input, (2) a glycan-based language model in the style of SweetTalk,^23^ and (3) a graph convolutional neural network based on SweetNet.^24^

**Figure 1.**
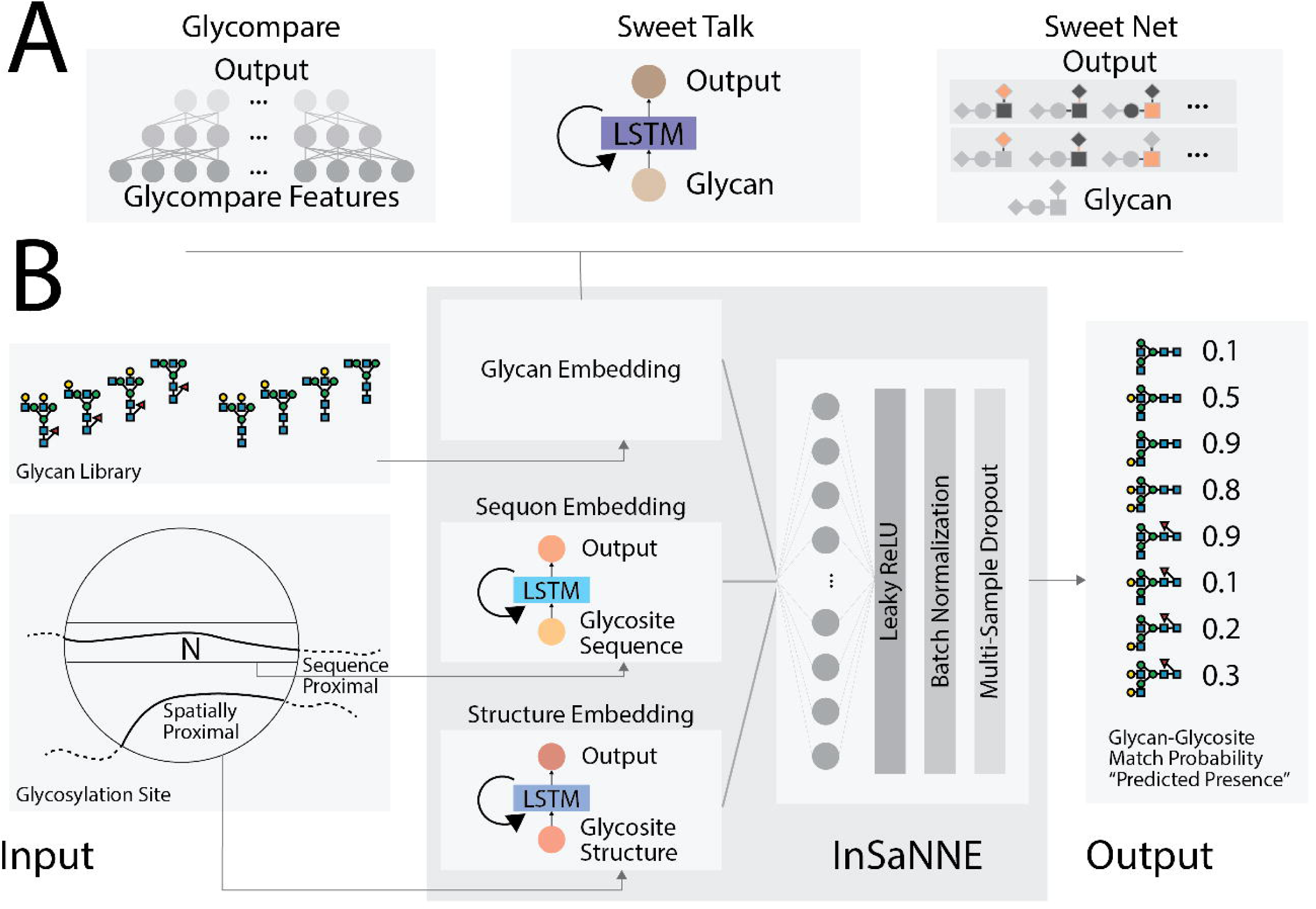
InSaNNE model architecture. **A)** Three model architectures were used to embed glycan structures in meaningful manifolds;^23,24,27^ given a glycan, these models output glycan-specific coordinates within the embeddings. To analyze the GlyCompare features of glycans, we used a fully connected neural network, while a SweetTalk-based language model used linear glycan sequences and a SweetNet-based graph convolutional neural network relied on glycan connectivity (see Methods for details). **B)** Full model architecture of InSaNNE. The results of one of the glycan embedding modules (A) is concatenated with protein-structure and protein-sequence embeddings output by the two protein-language models. These outputs were analyzed by a fully connected neural network and yielded the predicted probability of a glycan-glycosite match. Specifically, InSaNNE takes in a 14 amino acid glycosite-flanking sequence, optional spatially proximal amino acids, and a comprehensive library of 700 representative glycans on which InSaNNE was trained. Glycan libraries containing non-represented glycans can be used following additional training.

On average, the model based on GlyCompare glycan substructure features achieved a 76.3% accuracy in predicting which glycans have been observed at specific glycosites (**Table** 1). The recurrent neural network (SweetTalk; 79.9%) or graph convolutional neural network (SweetNet; 83.1%) models further improved the performance, demonstrating that optimizing the glycan analysis modules increases prediction performance. Choosing the SweetNet-based model as our best-in-class performer, we used stochastic weight averaging (SWA; Izmailov et al., 2019) to further optimize performance. SWA improved SweetNet-based model accuracy to 87.5% (**Table** 1) and was therefore selected as our final model and used for all downstream analyses.

**Table 1.**
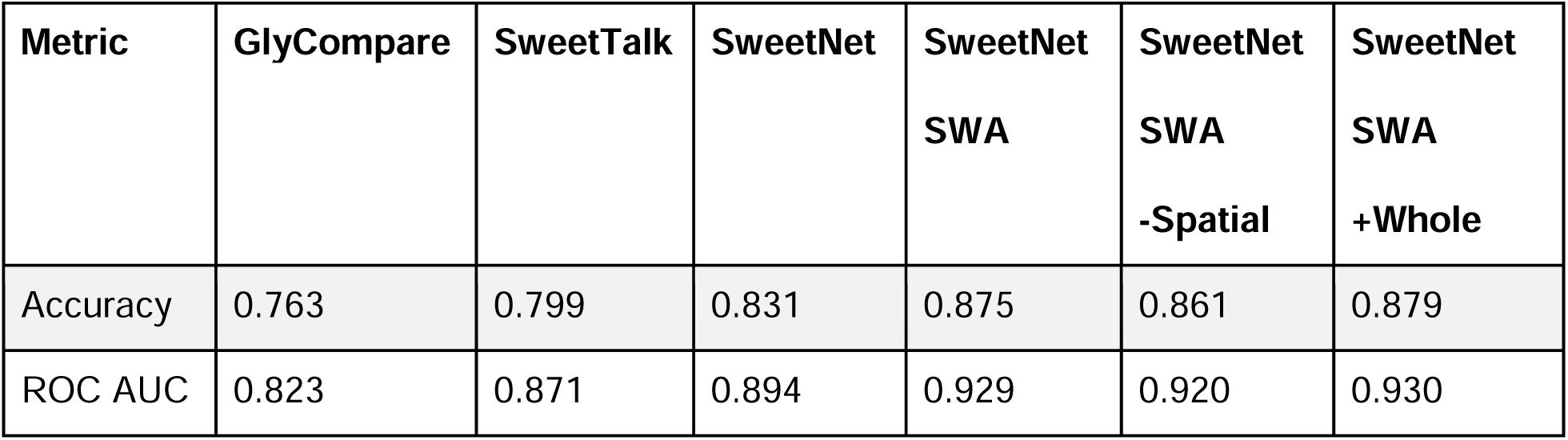
A model for glycan-glycosite matching was developed to predict permissible glycans on a glycosylation site. Modules analyzing the glycosite-flanking protein sequence and additional spatially proximal amino acids consisted of recurrent neural networks, while the module analyzing glycans was either a fully connected neural network using GlyCompare substructure features as input (GlyCompare), a glycan-based language model (SweetTalk), or a graph convolutional neural network (SweetNet). We further tested the effect of stochastic weight averaging (SWA) on model performance. Removing the information about spatially proximal amino acids from the model input is denoted by “-Spatial” while the addition of the whole protein sequence as an additional input for the model is indicated by “+Whole”. Results represent the mean values for accuracy and area under the curve (AUC) for the receiver-operator curve (ROC) on our test set after five independent training runs.

After optimizing the glycan analysis module, we analyzed the role of protein sequences on prediction performance. We trained a model that only had access to the glycan structure and the glycosite-flanking sequence, without additional spatially (3D) proximal amino acids. Compared to the full InSaNNE model (87.5% accuracy), the model without spatially proximal amino acids achieved a slightly worse performance (86.1%, **Table** 1). The marginal performance loss suggests that, while spatially proximal information helps, the glycosite-flanking residues are most important.

We next trained a model with access to the whole sequence of each protein, in addition to glycosite-proximal amino acids, and glycan structures. The additional information from the whole protein slowed training and inference, while providing a limited performance improvement (87.9% accuracy, **Table** 1). We concluded that distant amino acids carry limited relevant information for predicting permissible glycan structure that is not already captured in the nearby sequence and spatially proximal amino acids.

### Different glycosites prefer specific glycan features

True negatives, infeasible glycans, are hard to obtain experimentally, so we focused on recall (True Positive Rate). InSaNNE achieved a recall of 84.8% for *N*-linked glycosylation events in our dataset. The notable performance in these glycan-type-specific models, suggests that InSaNNE performs with exceptional recall – recovering most permissible glycans at a given glycosite.

Next, we examined which N-glycan motifs were more difficult for InSaNNE to predict. For this, we calculated the average prediction accuracy for each glycan feature in the validation set. Several rare glycan motifs (<10 observations) were more difficult to predict (**Figure 2**a). However, InSaNNE exhibited a predictive accuracy of >80% for most motifs (**Figure 2**b). Since glycan features represent a hierarchical feature set, rare motifs with low prediction accuracy are not independent from each other and formed clusters based on glycan structure similarity (**Figure 2**c). For example, glycan features with lower predictive performance were enriched for oligomannose. Analogous to the glycan features, most glycosites exhibited an aggregate predictive accuracy >90% (**Figure 2**d) and we found prediction performance correlated with the number of observed glycans for similar glycosites (close in the embedding manifold; **Supplementary Figure 1**). Predictions were robust to the removal of single amino acids or short motifs, suggesting redundancy within glycosite-flanking sequences and soft boundaries on the flanking window size (**Supplementary Figure 2**). Furthermore, the flanking residues, rather than the central sequon-proximal residues, informed model predictions the most; ablation of upstream residues was most impactful on performance (**Supplementary Figure 2**). In general, given the consensus sequence of *N*-linked glycosylation, flanking residues are more variable, and may carry more information for deep learning models, than more conserved sequon-adjacent residues.

**Figure 2.**
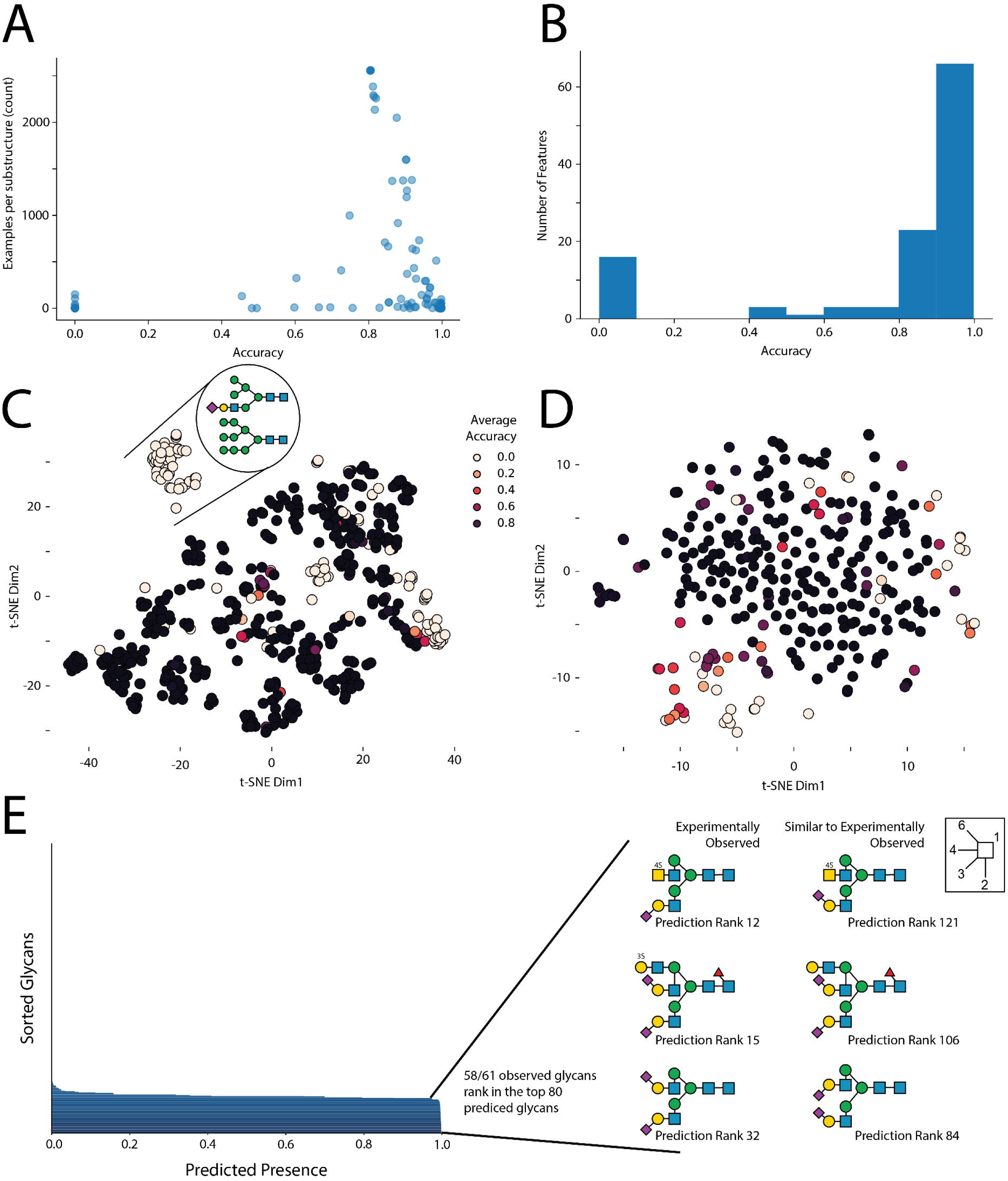
Characterizing the glycan-glycosite-matching model InSaNNE. **A)** Dependence of glycan feature prediction performance on occurrence. Using our trained InSaNNE model, we plotted the averaged prediction performance of glycan features against their counts in our dataset. **B)** Glycan feature accuracy distribution. A histogram of the prediction performance for all observed glycan features is shown. **C)** Clusters of difficult-to-predict glycan features. We used t-SNE to visualize the glycan representation learned by InSaNNE for all glycan features. Each feature was colored by its averaged prediction performance to identify structurally related clusters of glycan features that are more difficult to predict for InSaNNE (shown in brighter colors). **D)** Prediction performance depending on the glycosite was visualized using a t-SNE of the glycosite representations learned by InSaNNE. For all glycosites in our dataset, we averaged prediction performance over all glycans and colored glycosites by prediction performance to identify difficult to predict glycosite clusters. **E)** Experimentally observed and predicted glycans at a glycosylation site of human uromodulin were compared. GTVLTR<u>N</u>ETHATYS (P07911:N396) was used to predict permissible glycans using the trained InSaNNE model, and the top 80 predicted glycans were analyzed and compared to previously observed glycans at that site ^32^.

To illustrate the capabilities of InSaNNE, we used the model to predict the feasibility of all glycans in our dataset at the glycosite GTVLTR<u>N</u>ETHATYS (P07911:N396) from human uromodulin – the most abundant protein in human urine and relevant for chronic kidney disease.^32^ Notably, 58 of 61 experimentally observed glycans were placed in the top 80 predicted glycans (**Figure 2**e). Additionally, top glycans that were not previously reported at this glycosite shared features with the observed glycans, such as a strong negative charge via sialylation and/or sulfation. These results further demonstrate protein-sequence-based glycan prediction and emphasize the value and relevance of our model.

### Single amino acid changes modulate specific glycan features

While the ablation of individual glycosite-flanking amino acids does not substantially diminish model performance (**Supplementary Figure 2**), glycosylation efficiency and range can be impacted by glycosite-flanking mutations.^5,6,9,11^ Therefore, we tested if InSaNNE can predict how changes to the glycosite-flanking sequence will impact glycosylation. This could facilitate glycoengineering and elucidate structural interactions between protein and glycan structures at the glycosylation site. We performed a deep mutational scan *in silico* (replacing each of the 14 glycosite-flanking amino acids with all amino acids) on every N-glycosite in our dataset. Using the modified glycosite sequences as inputs for InSaNNE, we analyzed the changes in predicted glycans compared to the wild-type sequence. To focus interpretation, we grouped glycans into “sialylated” and “fucosylated.” This allowed us to track the changes in predicted probability for each of these features following specific glycosite-flanking mutation (**Figure 3**, **Supplementary Figure 3**). However, while these reflect general trends of individual glycosites across all proteins, amino acid substitutions may have effects that deviate from these general trends.

**Figure 3.**
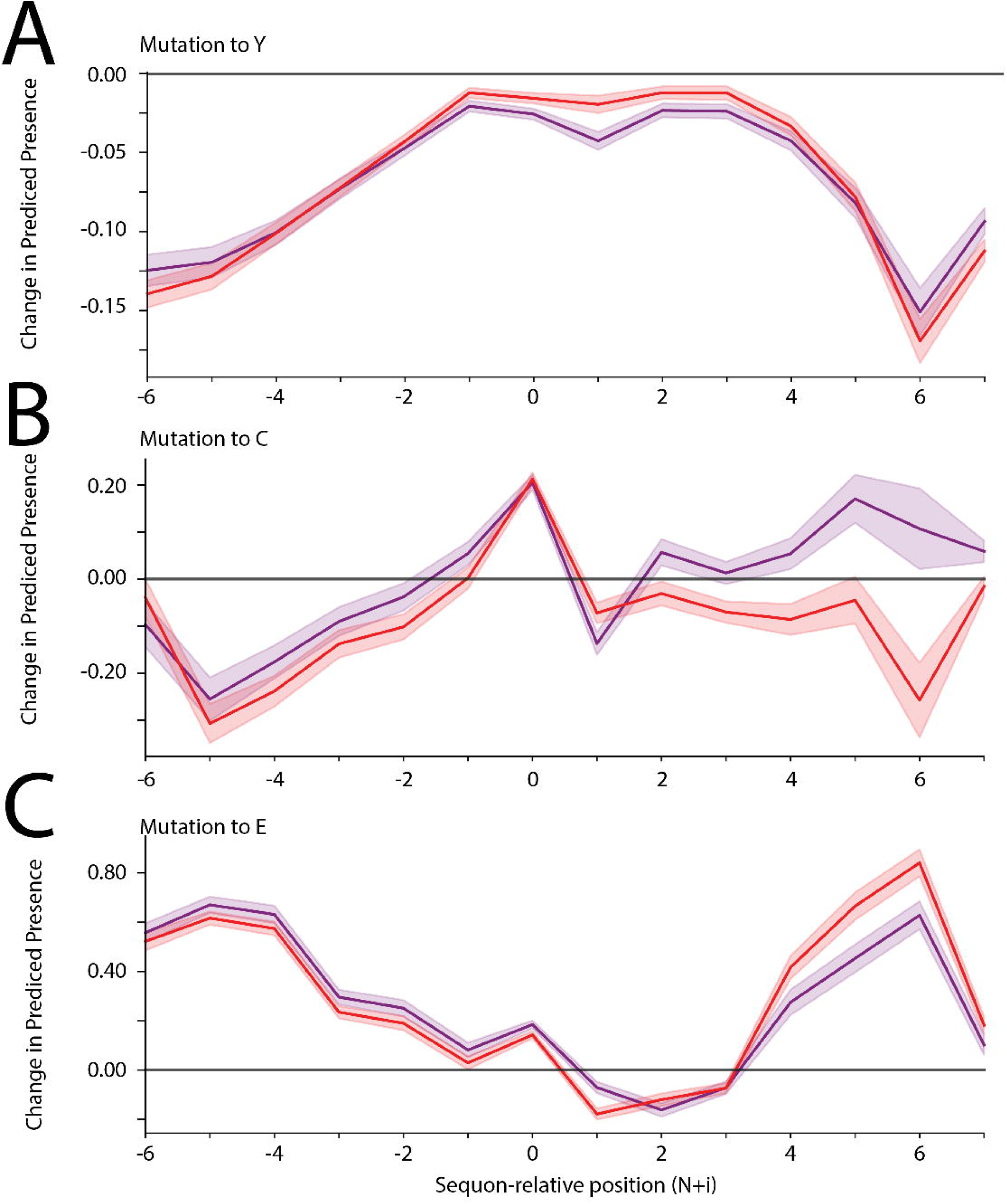
Effects of amino acid substitutions on predicted glycosylation ranges. A-C) For all N-linked glycosites in our dataset, we substituted each amino acid with tyrosine (A), cysteine (B), or glutamate (C) and input the modified glycosite-flanking sequences into our InSaNNE model and predicted feasible glycosylation. We then calculated the average change (predicted presence difference) compared to the predicted wild-type glycosylation glycosites; shown here with a 95% confidence interval. Lines for changes to fucosylated (red) and sialylated (purple) glycans are shown. See **Supplementary Figure 3** for analogous plots for other amino acid substitutions.

For multiple amino acid substitutions, we observed distinct changes in the predicted glycosylation of modified glycosites, with clear differences between changes to upstream and downstream regions. The introduction of some amino acids (e.g., tyrosine; **Figure 3**a) had the same qualitative effect regardless of where they were introduced. Meanwhile, other amino acids (e.g., cysteine; **Figure 3**b) have diverging effects, with a decrease in predicted complex glycans when introduced upstream and an increase when it is present downstream. We also observed that predicted changes in glycosylation were impacted more strongly by mutations in the distal parts of the glycosite-flanking sequence (e.g., glutamate; **Figure 3**c). These general trends of amino acid-glycan associations could be useful for glycosite-specific glycoengineering.

### Uncharacterized glycoproteins and glycan compositions can be annotated with candidate glycan structures

Computational prediction to annotate protein features and functions is done routinely for newly discovered proteins, yet limited *in silico* characterizations exist for glycosylation. However, the relative speed of predicting glycosylation would make it invaluable for new, existing, or poorly characterized proteins; typical glycoprofiling approaches can otherwise take several months. Even many well-characterized glycoproteins have only compositional measurements (unstructured monosaccharide counts) since glycan structure measurement and characterization are resource and expertise-intensive processes. Thus, InSaNNE could be invaluable for annotating glycosylation sites.

Predicting glycosite location is one of the few high-confidence bioinformatic predictions involving glycosylation.^33–37^ To extend this capability, we predict the feasible glycan structures of 2,763 human N-linked glycosites in the GlyConnect database.^30^ For this, we used InSaNNE to analyze the annotated glycosylation sites together with the six upstream and seven downstream amino acids. For each glycosite, we predicted the likelihood of 199 N-linked glycans (**Supplementary Dataset 1**). Using our independent test set, we ascertain a threshold with an acceptable false-positive rate (AUC 0.92, **Figure 4**a). A threshold of 0.6 (predicted presence) corresponded to a false-positive rate <10% while maintaining a true positive rate >85%. This allowed us to assess the recall or sensitivity of our predictions within GlyConnect by quantifying known glycan structures that were successfully predicted (**Figure 4**b). Thus, InSaNNE could inform future experiments and comparative analyses of structure-based constraints in glycosylation and functional impacts.

**Figure 4.**
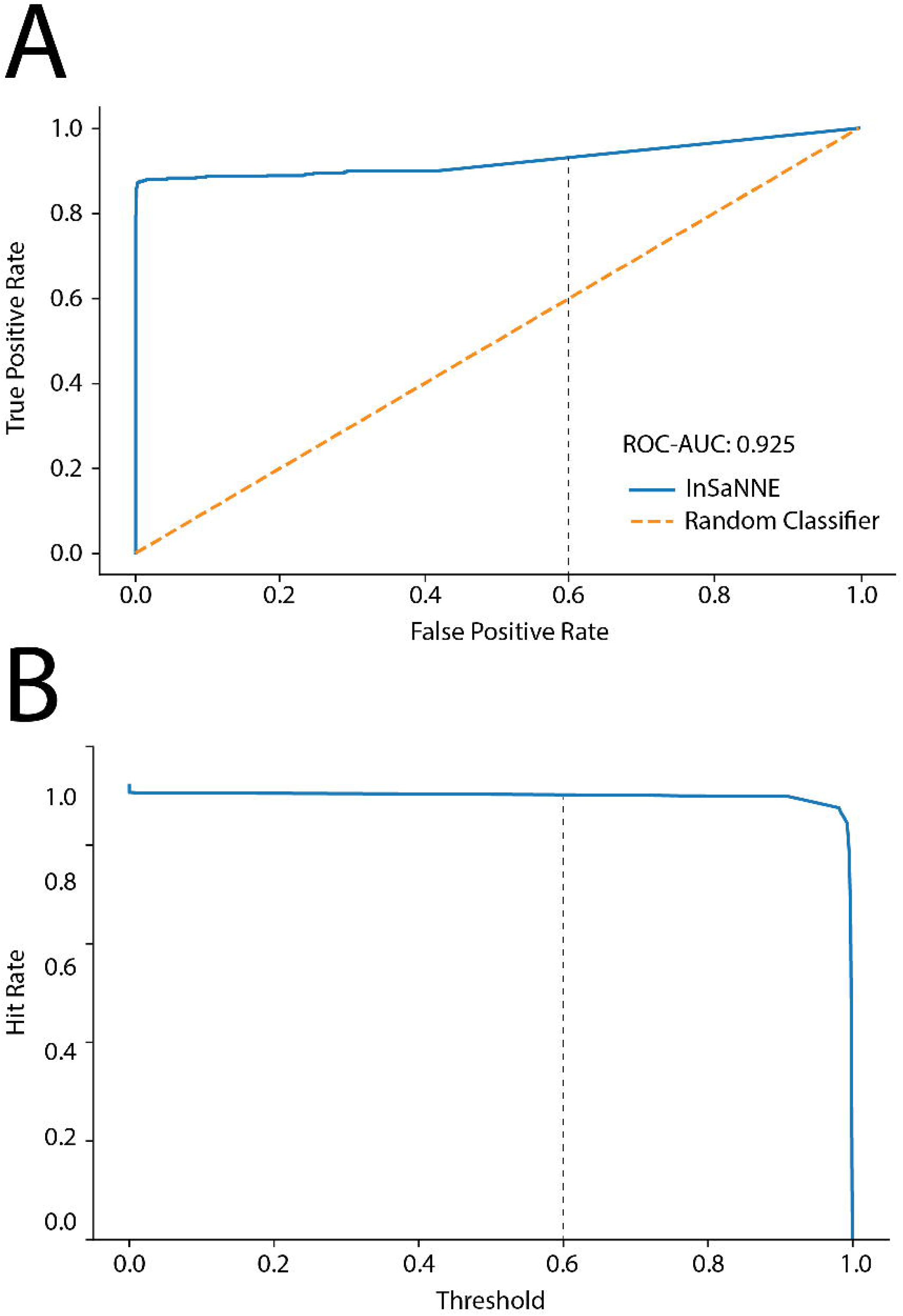
Enriching GlyConnect with InSaNNE predictions. **A)** For classification thresholds between 0 and 1, we assessed true and false positive rates of InSaNNE predictions on the independent test set and compared it to a random classifier baseline. **B)** We validated InSaNNE predictions with existing structures on GlyConnect by investigating the influence of classification threshold on the hit rate (i.e., recall/sensitivity) of InSaNNE accurately predicting known glycan structures in GlyConnect. The grey dotted line marks the 0.6 threshold used.

### InSaNNE predicts complex glycans in the enhanced aromatic sequon and the SARS-CoV-2 Spike

N-glycans are commonly grouped into categories, such as highly processed complex glycans, hybrid glycans, and immature oligomannose glycans.^38^ Previous work showed that an aromatic residue located two-positions N-terminal from a glycosylation site results in less complex N-glycosylation at the site, termed the enhanced aromatic sequon.^6^ In this case, an L to F substitution two residues upstream of the CD2 glycosylation site transformed the site from predominantly complex (sialylated) and hybrid structures to low complexity (oligomannose) structures. When InSaNNE evaluates the same sequences, the F allele sequence shows significantly higher predicted presence for higher-mannose structures. We predict an enrichment for 7-mannose structures (One-sided Mann-Whitney-Wilcoxon, p=0.017) and predict an overall increase in oligomannose structure for the F allele (Linear model; Wald, p<0.001; F-statistic, p=7.44×10^−5^; **Figure** 5**a**). We see a corresponding decrease in sialylated structures in the F allele (One-sided Mann-Whitney-Wilcoxon, p<1e-4; **Figure** 5**b**).

**Figure 5.**
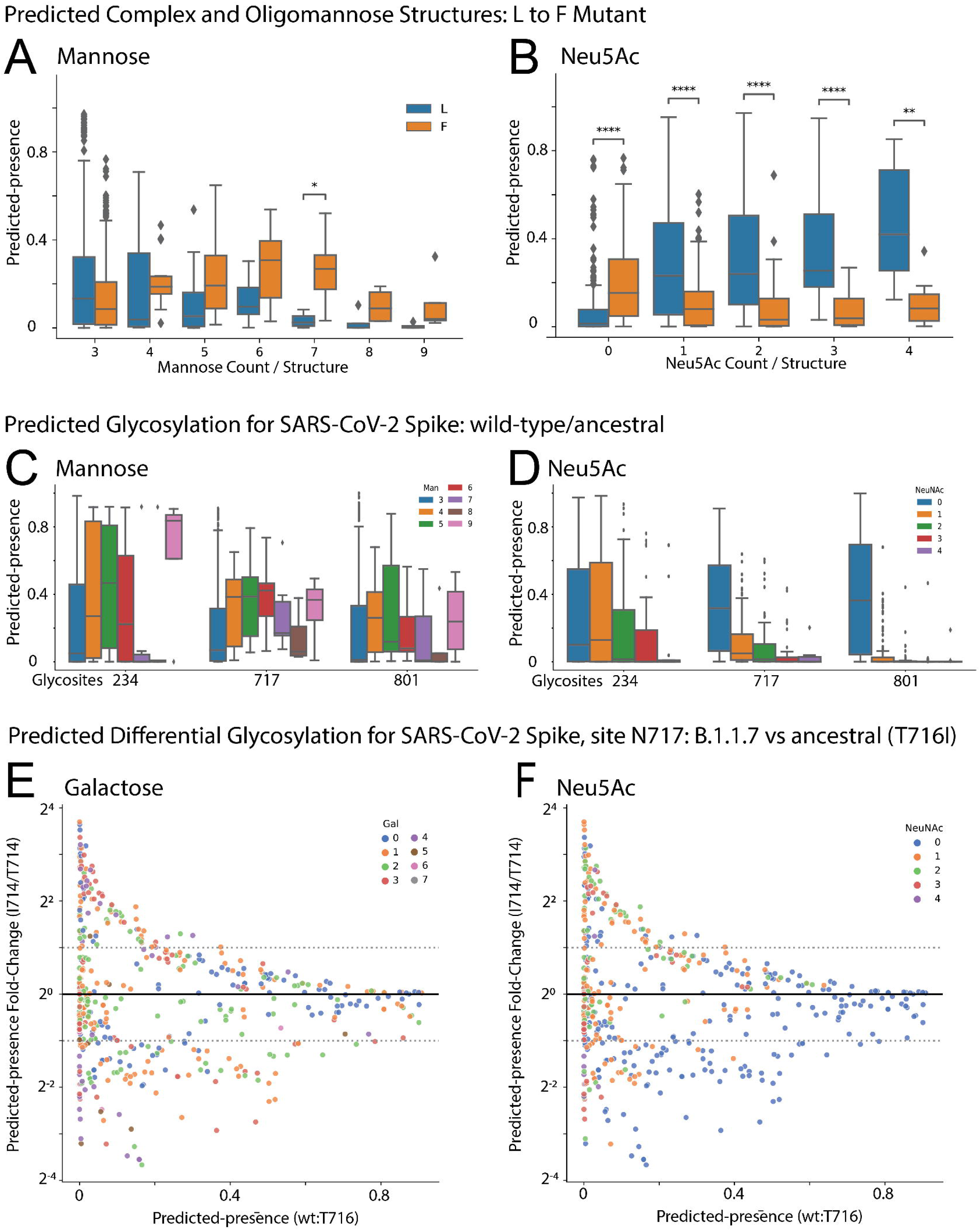
InSaNNE predicts complex glycans around the enhanced aromatic sequon and the SARS-CoV-2 spike protein. (A-B) Boxplot distributions of predicted-presence for the L and F variants at N-2 stratified by number of **(A)** mannoses per glycan and **(B)** sialic acids per glycan. **(C-D)** Boxplots describing predicted glycosylation by (C) mannose per glycan and (D) sialic acid per glycan for three oligomannose sites in the SARS-CoV-2 spike glycoprotein. See **Supplementary Figure 4** for all SARS-CoV-2 spike glycosylation sites. **(E-F)** Fold changes of predicted glycans at site N717, labeled by number of (E) galactose and (F) sialic acid units, between the wild-type and B.1.1.7 spike protein. Predicted-presence fold-change (y-axis) is stratified by the basal predicted-presence for each glycan in the wild-type (x-axis). Predicted-presence fold-change from wild-type by galactose, mannose, GlcNAc, and sialic acid is provided for N717 and N616 in B.1.1.7 (**Supplementary Figure 5**) and D615G **(Supplementary Figure 6)** variants respectively. ns: p>0.05, *: p<0.05, **: p<0.01, **: p<0.001, ***: p<1e-3, ****:p<1e-4

InSaNNE also recapitulates glycan types of SARS-CoV-2. These sites have been extensively characterized throughout the pandemic.^15,39–41^ N234, N717, and N801 are highly reproducible oligomannose sites.^15^ Oligomannose at N234 is consistently high (80-100%)^15^ and appears necessary to support the open ACE2-binding spike conformation.^42^ Our predictions show strong preference for Man5 and Man9 structures and a strong anticorrelation with sialylation (**Figure** 5**c-d**). Sites N717 and N801^15^) are predicted here to have almost no sialylation (**Figure** 5**c-d**). Predictions for all glycosylation sites were mostly consistent with empirical observations (**Supplementary Figure 4**).

We wondered if the spike protein of new strains shows predictable changes in glycosylation. We examined InSaNNE predictions at site N616 in a simulated D614G variant (**Supplementary Figure 6**) and N717 in a T716I variant (**Figure** 5**e-f**). We found distinct changes in predicted glycosylation. T716I, between the furin cleavage site and the fusion peptide, is within the more conserved S2 sequence and retains moderate antibody accessibility regardless of RBD conformation.^43^ To focus on relevant changes, we examined those with non-negligible ancestral predicted-presence (>0.1) and substantial fold change (|logFC|>1) relative to the ancestral spike. At site N717 in the T716I variant, many asialylated sugars with one to three galactose residues decrease relative to ancestral (**Figure 5f**, blue points). Additionally, a small number of sugars with zero to two sialic acids and one to four galactose residues increase. Though InSaNNE predicts that site N717 becomes variably permissible to mono-, di-, tri- and tetra-antennary sialylated and asialylated structures, empirically, it is an oligomannose site, suggesting these terminal galactoses may not be visible without additional mutations to the site. Distinctly, InSaNNE reveals few confident changes at site N616 in the D614G variant (**Supplementary Figure 6**). If glycan structure can be predicted from primary sequence, site occupancy may also be bound by these constraints.

### InSaNNE predictions recapitulate biantennary abundance on human IgG3

Mutations can perturb glycosylation in IgG3.^9^ Eight complex biantennary structures in human IgG3 were measured for wildtype (*wt)* and glycosite (N297; P01860:N227) proximal mutants. While the *wt* IgG3 showed a preference for core-fucose and a1-6-branch galactose, R301A increased all terminal galactose, and Y296A accepted no galactosylation (**Figure** 6**a**). Thus, primary protein structure can profoundly influence glycosylation.

**Figure 6.**
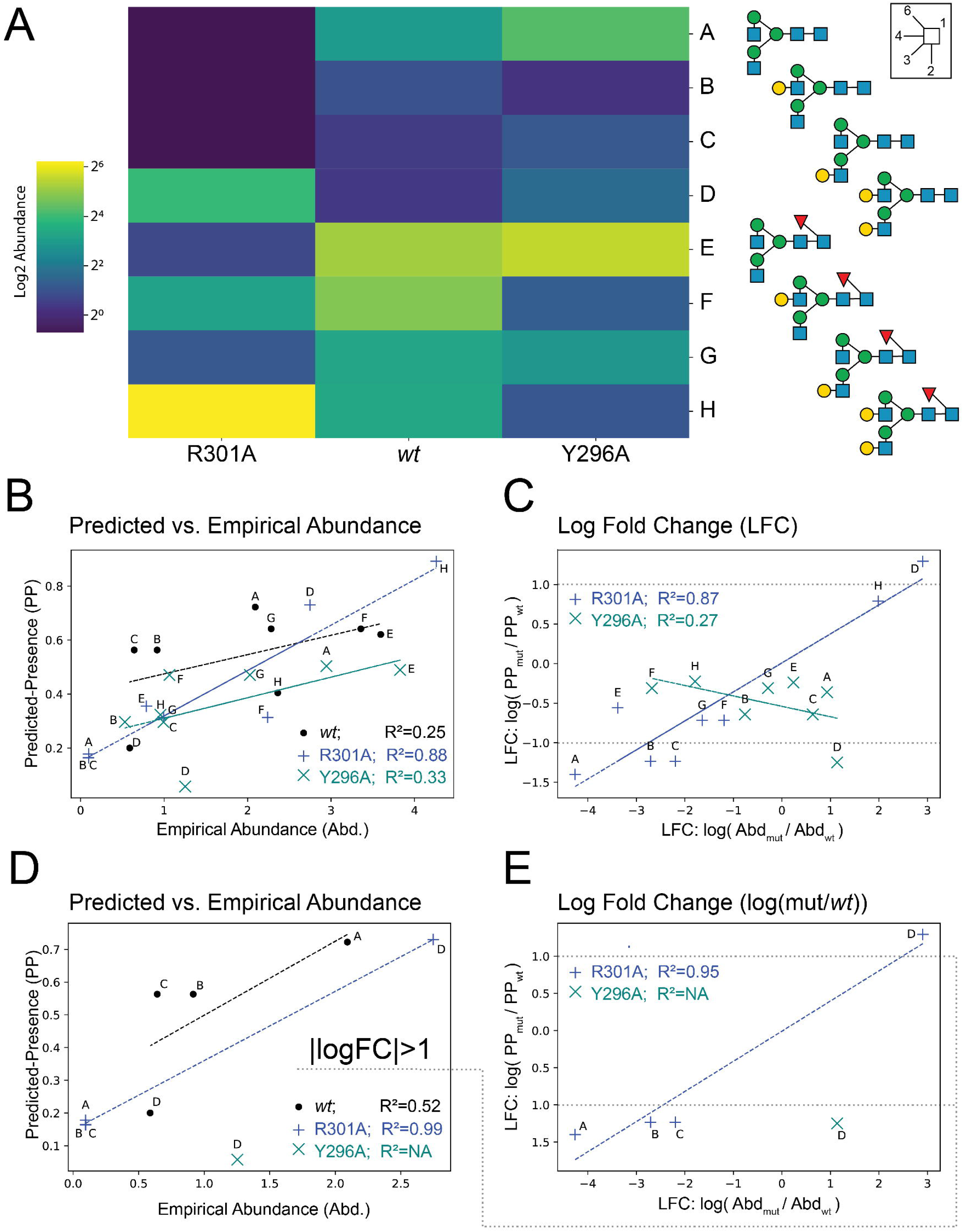
InSaNNE predictions of relative abundance on IgG3. **A)** Heatmap showing the log-scale abundance of various glycan species observed in wt and mutant Fc on human IgG3.^9^ **B)** The background-adjusted InSaNNE predicted-presence is compared with the empirical abundance in wild type (black), R301A mutant (blue), and the Y296A mutant (teal). **C)** Log fold change between glycan abundance for mutants relative to wildtype were compared between empirical and predicted abundance for all glycans. **D-E)** The bottom panels mirror panels **B-C** except glycans with a predicted absolute log fold-change less than 1 were removed.

We compared InSaNNE predictions for the R301A and Y296A mutants and found that predicted-presence and change in predicted-presence were correlated with empirical occupancy. Abundance-prediction correlation was high for the R301A mutant (R^2^=0.876; **Figure** 6**b**) and moderate for *wt* abundance (R^2^=0.25; **Figure** 6**b**). Predicted presence was consistent with measured abundance in the Y296A mutant (R^2^=0.33; **Figure** 6**b**). Interestingly, prediction performance increased when we compared changes relative to *wt*. The predicted presence log fold-change in R301A relative to *wt* was highly correlated with measured abundance log fold-change (R^2^=0.87; **Figure** 6**c**). Yet, the consistency in predicted vs observed change for Y296A decreased dramatically (R<0, R^2^=0.27; **Figure** 6**c**). To further probe the prediction failure in Y296A, we removed glycans with small predicted changes (|logFC|<1). Without the low-confidence changes, abundance prediction performance for *wt* (R^2^=0.52), R301A (R^2^=0.99), and log fold-change (R301A vs. *wt*: R^2^=0.95) improved (**Figure** 6**d-e**), while nearly all predictions for Y296A dropped out. These results suggest that InSaNNE can predict occupancy and occupancy change for non-small (|log fold-change|>1) changes.

### Accessing InSaNNE predictions and continuous comparison through GlyConnect

We evaluated the agreement between InSaNNE predictions and GlyConnect data at the compositional level. **Figure** 7**a** shows the protein-page d3 heatmap illustration comparing GlyConnect-annotated glycosylation events for human coagulation factor XI (UniProt:P03951; GlyConnect:818) with InSaNNE predictions; GlyConnect:818 is supported by four published references. **Table 2** summarizes the comparison between GlyConnect annotation and InSaNNE predictions for human coagulation factor XI.

**Figure 7.**
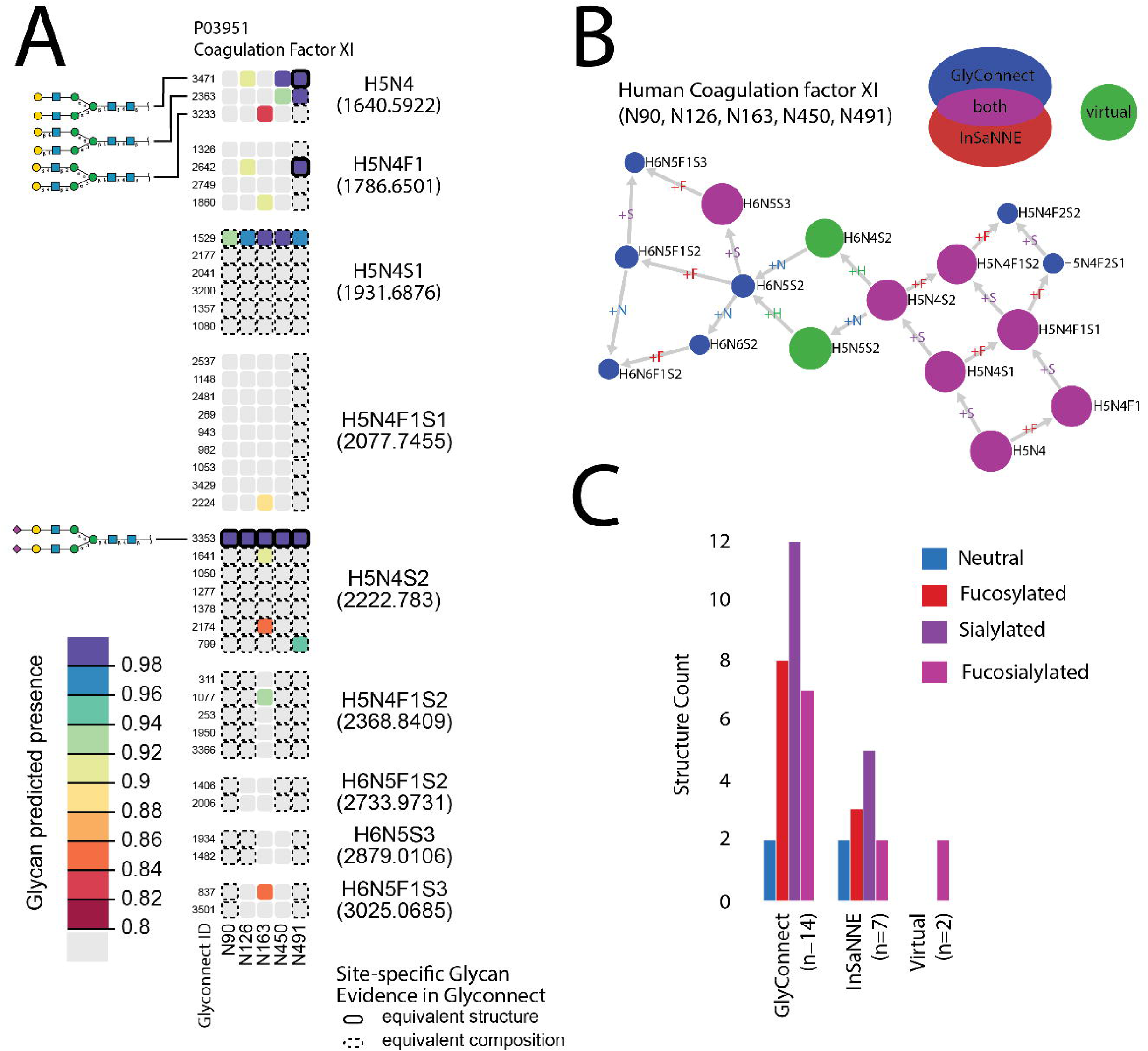
Predicted glycosylation pattern of human coagulation factor XI (P03951). H: hexose, N: hexosamine, F: fucose, S: sialic acid. (**A**) The heatmap displays the predicted presence for glycan structures at each known N-glycosite and indicates agreement with glycans previously observed at those sites retrieved from GlyConnect. The structures in each row are ordered by glycan composition; columns represent the five annotated N-glycosites of P03951. Site-specific glycan structure predictions are many-to-many relationships in the GlyConnect database since the same structure may be associated with several sites and conversely a single site may be predicted to present several similar yet non-mutually exclusive glycan structures. Composition blocks contain all structures matching a specific composition. Color indicates the strength of the predicted presence from 0.8 (lower-bound cutoff) to 1 (predicted presence upper-bound). A solid-line borders indicate exact structural matches (identical precise monosaccharides and identical linkages) while dashed lines indicate composition matches (monosaccharide category, e.g., hexose) with at least one non-identical linkage; composition-equivalent blocks (e.g., H5N4) are labelled. (**B**) A Compozitor graph representing compositional similarity between predicted and observed glycans. Fourteen glycan compositions are reported in GlyConnect for human coagulation factor XI. Nodes are connected via single monosaccharide additions represented as the edge label. Seven compositions are predicted and all included in the fourteen previously observed compositions (magenta). Two virtual nodes (green) were added to connect the graph. Numbers within the blue nodes express a correspondence in GlyConnect data between a composition and structures. When the number is absent it means we only have compositional data. The size of the non-blue nodes represents a comparison with the total content of GlyConnect to indicate the likelihood of the composition. For large nodes, the composition occurs often, irrespective of the protein where it is seen. (**C**) The bar chart represents glycan properties mapped in all subsets (database, predicted and virtual). It highlights the similarity across properties of predicted and stored structures.

**Table 2.**
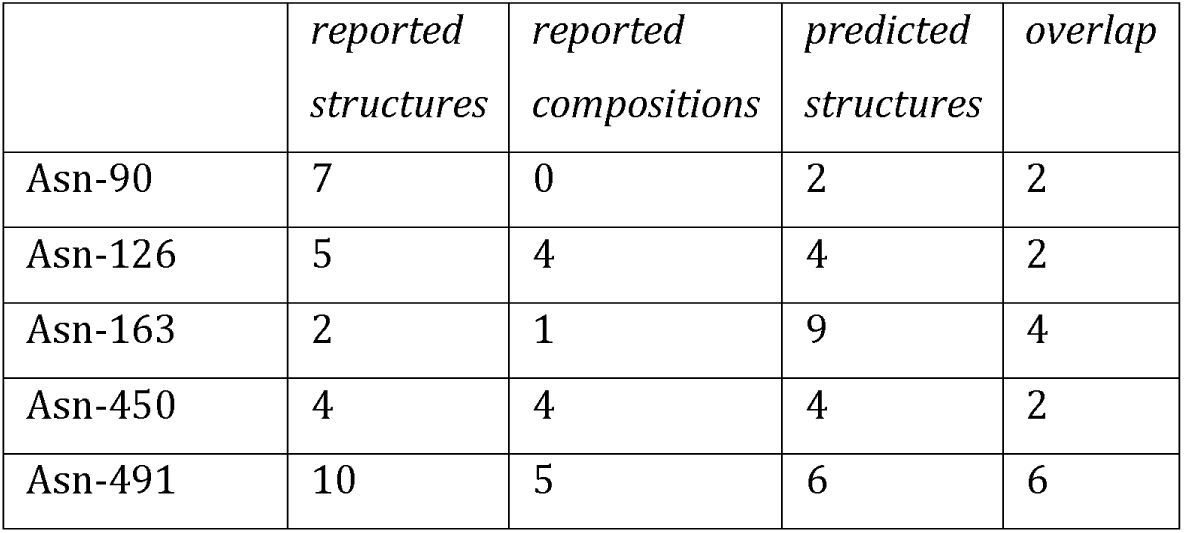
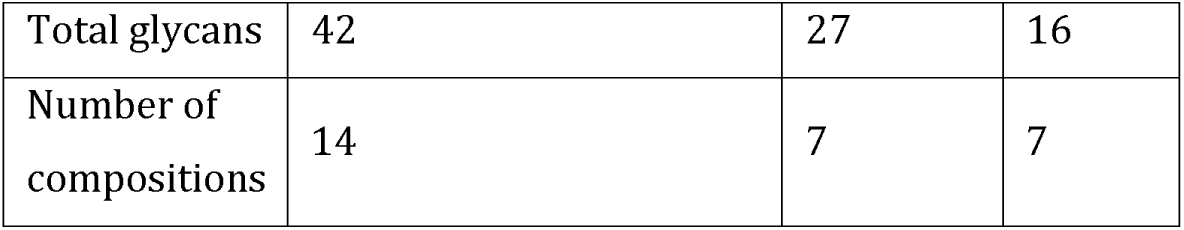
Summary of knowledge of human coagulation factor XI (P03951) as stored in the GlyConnect database at structural (first column) and compositional (second column) resolutions along with predicted structures (third column). The overlap between stored and predicted (predicted presence ≥ 0.8) structures is shown in the fourth column and the last row features the overall number of compositions. Note that overlap refers to matches between predicted structures and reported structures or compositions; one reported composition can map to multiple predicted structures.

|  | *reported*<br>*structures* | *reported*<br>*compositions* | *predicted*<br>*structures* | *overlap* |
| --- | --- | --- | --- | --- |
| Asn-90 | 7 | 0 | 2 | 2 |
| Asn-126 | 5 | 4 | 4 | 2 |
| Asn-163 | 2 | 1 | 9 | 4 |
| Asn-450 | 4 | 4 | 4 | 2 |
| Asn-491 | 10 | 5 | 6 | 6 |

*Table 2 - Summary of knowledge of human coagulation factor XI (P03951) as stored in the GlyConnect database at structural (first column) and compositional (second column) resolutions along with predicted structures (third column).
|  |  |  |  |
| --- | --- | --- | --- |
| Total glycans | 42 | 27 | 16 |
| Number of compositions | 14 | 7 | 7 |

The first composition, H5N4 (five hexoses and four hexosamines), matches three structures with similar linkages recorded in GlyConnect. At site P03951:N491, in composition block H5N4, we see InSaNNE correctly predicts the presence of GlyConnect glycan 3471; the dashed-line compositional matches to glycans 2363 and 3233 are expected as all three glycans are members of the same composition block. Additionally, glycan 2363 is highly predicted at N491 suggesting a partial linkage resolution for the incompletely determined structure stored in GlyConnect. Likewise, structures matching the H5N4S2 (five hexoses, four hexosamines, and two sialic acids) compositions contain glycan 3353 predicted and observed at all sites. Within composition block H5N4S2, InSaNNE predicts a higher likelihood (>0.9) for glycan 1641 at N163 (biantennary α2,3-Neu5Ac). Glycan 1641 offers a complete resolution of structural ambiguity for H5N4S2 at N163. Prediction and annotation both involve flexible linkage definitions, particularly for non-core residues. In contrast, the prediction at site N163 is more extensive than reported data. Interestingly, N163 is a rare NXC sequon, which may explain the smaller number of reported structures and provides novel insights into the distinct preferences of this rare sequon.

For human coagulation factor XI, GlyConnect contains site-specific observations of 42 structures and compositions, and 14 additional distinct but structurally related glycans (Table 2). Compositional similarity was displayed using Compozitor (**Figure** 7**b**). The Compozitor graph shows 14 compositional nodes connected through the addition of a single monosaccharide. Two virtual nodes (green: H6N4S2 and H5N5S2) are needed to fully connect the graph.^44^ All site-specific InSaNNe-predicted structures correspond to previously annotated site-specific compositions in GlyConnect (magenta). InSaNNE fails to predict structures corresponding to three previously reported compositions the H6N5S2, H6N5F1S2, and H6N5F1S23. Interestingly, the glycan property distribution (**Figure** 7**c**) is similar between reported and predicted compositions, suggesting a lack of systematic bias that would diminish expected performance for specific glycotypes. Other compositions were found in large scale glycoproteomics experiments without any precise structural features and may be less reliable annotations.

## Discussion

Here we present InSaNNE, the Interloping Saccharide Neural Network Extrapolation, for predicting glycans on membrane-bound and secreted proteins. This approach employs a recurrent neural network and a graph convolutional neural network with stochastic weight averaging to predict feasible glycan structures based on the underlying protein sequence. InSaNNE successfully predicts known glycan structures on a wide range of proteins and assesses the impact of single amino acid substitutions on resulting glycan structures. Beyond initial cross-validation and test-set validation, we successfully predicted glycans on uromodulin, SARS-CoV2, IgG3, and across the GlyConnect database. We have added the glycan predictions to the glycome database GlyConnect, making them accessible for further study of this discovery. Importantly, InSaNNE further questions the premise of template-free glycan biosynthesis. Glycosylation through the bounded biosynthesis paradigm, and its accessibility through the InSaNNE framework, will facilitate more accurate and accessible study of diverse glycoproteins and glycoproteomic behaviors.

InSaNNE enables the draft annotation of glycosylation on novel proteins, glycoprotein composition analyses, glycoinformatics, and whole proteomes. By increasing the predictability of glycans, we have reduced the challenge of measuring glycans. Mass spectrometry is the gold standard in glycan measurement today, but these measurements may produce partially ambiguous structures and topologies. Consequently, the field is rich with datasets and databases of partially or minimally assembled glycoprofiles.^45–48^ Combining measured glycan compositions with site-specific predictions of feasible glycosylation should facilitate automated glycoprofile assembly. These annotations can be completed for novel and existing glycoprofile assemblies; because of the automated nature, structural glycoprofiles can be assembled for single experiments or entire databases with comparable ease. The sequence-only nature of the prediction is especially important, as many proteins lack experimental structural observations; an algorithm that can operate on the primary sequence is considerably more portable than one requiring structural information. A sequence-only prediction can even be used to quickly compare different isoforms or predict glycans on newly discovered protein sequences.

We demonstrated our ability to glycosylate an entire proteome by predicting decoration throughout GlyConnect. Newly glycosylated proteins can be used to identify lectin-binding, glycan co-ligands, alternative charge, or steric conformations on proteins of interest, and changes in protein dynamics. These predictions can be disseminated to enrich databases detailing glycosylation^30,49,50^ and other post-translational modifications,^51–53^ protein structure,^54,55^ domains,^56,57^ and interactions.^58–61^ Future work will extend this approach to *O*-linked glycans, an even more challenging endeavor due to less available data for training and a seeming absence of a clear consensus sequence on the protein side.^62^

Predicted glycosylation can be used to inform large genetic and genome-wide studies. Genetic variation can change protein function and resulting phenotype, but here we demonstrate that it can impact glycosylation. InSaNNE can predict such changes and thus provide further hypotheses for elucidating disease mechanisms. For example, adding predicted differential glycosylation to a study of a high-heterogeneity critical immune gene like Human Leukocyte Antigen (HLA) will be invaluable. This is because HLA has a functional binding-groove adjacent glycosite^63,64^ that could contribute to the behavior, accessibility, and peptide presentation. Some HLA molecules have already been observed to carry allotype-specific glycans.^65^ Beyond HLA, understanding differential glycosylation on reference and variant molecules can help distinguish benign from pathogenic mutations: characterized (e.g., ClinVar) or uncharacterized (e.g., precision medicine). Additionally, certain glycoforms can modulate secretion.^66,67^ Because each glycan may confer a change in behavior, phenotypes of highly diverse glycoproteins such as secretion, protein-ligand interactions, cell-cell interactions, and extracellular protein complexes can be enriched by knowledge of glycosylation. These are only a few of the studies that may benefit from protein-predicted glycosylation potential.

Bounded biosynthesis provides a more complete picture of immune evasion by evolving pathogens. Glycan-coated viruses have been responsible for many pandemics, while nearly every decade has seen epidemic strains of viruses, such as influenza. Recent work has highlighted the alignment of these fluctuations with changes in glycans decorating these viruses.^12^ Without specific glycoforms, it is not possible to determine which of these viruses successfully disguised critical immune epitopes and which viruses created or maintained new lectin-targeted epitopes. With specific glycan prediction, we may predict the most concerning mutations, those that may reinforce a glycan shield,^11,68–70^ stabilize virulence factors,^42^ or occlude immunogenic antigens.^71^ Glycoform predictions can provide these missing data along with previously inaccessible insight into the history and future of viral evolution.

In summary, bounded glycan biosynthesis, as functionalized by InSaNNE and made accessible through GlyConnect, will enable investigators to easily consider glycosylation across many areas of biological study. InSaNNE will thereby sharpen our understanding of the extracellular space and innumerable intercellular phenotypes.

## Supporting information

Supplemental Marterials

## Acknowledgements

This work was supported by NIGMS (R35 GM119850, NEL), the Novo Nordisk Foundation (NNF20SA0066621, NEL), and a Branco Weiss Fellowship – Society in Science awarded to D.B.

## Conflicts

This work is associated with a provisional patent filed by the authors, and Augment Biologics, founded by BK and NEL.

## Methods

### Site-specific glycosylation training set construction

Empirical site-specific glycosylation data from humans was obtained from UnicarbKB^29^ and Glyconnect^72^ with supplemental information from GlyGen.^73^ The protein structure annotation was done using the Structural Systems Biology (ssbio) package in python.^74^ Protein structure analysis was performed in Python v2.7.15 using ssbio v0.9.9.8 to retrieve and calculate: existing empirical and homology models from PDB and SWISSMOD (PDBe SIFTS),^75^ *de novo* homology models (I-TASSER v5.1), sequence properties (EMBOS v6.6.0.0 pepstats), sequence alignment (EMBOS v6.6.0.0 needle), secondary structure (DSSP v3.0.0, SCRATCHv1.1::sspro and SCRATCHv1.1::sspro8), solvent accessibility (DSSPv3.0.0 and FreeSASAv2.0.2), and residue depth (MSMSv2.2.6.1). Additional amino acid aggregate features were calculated using R::seqinr. Glycan structures were annotated using a combination of glypy^76^ and GlyCompare^27^ for structure parsing and comparison, respectively. All glycan substructures, a connected subset of monosaccharides with and without linkage information, were extracted from each glycan, merged to make a superset of substructures, then mapped to each glycan. This resulted in a mapping from every glycan in the input database to shared substructures.

For the dataset used to train InSaNNE, we extracted 1,721 unique glycosylation events from UniCarbKB.^29^ This included the glycan structure that was observed and the glycosite-flanking sequence (14 amino acids, with the glycosylated amino acid in the center) and structural information in the form of additional amino acids within 6Å if structural simulations converged. As negative examples, we generated the same number of combinations of glycosites and glycans that have not been observed.

### Model construction

All glycan-glycosite matching models comprised (1) a recurrent neural network that analyzed the amino acid sequence of the glycosite, (2) another recurrent neural network analyzing the amino acids of the three-dimensional glycosite surroundings, (3) a model analyzing the glycan structure, described below, and (4) a part consisting of fully connected layers to use the concatenated features generated by the previous modules to predict whether a glycan is permissible at a glycosite. The recurrent neural networks consisted of a 128-dimensional embedding layer followed by two bidirectional long short-term memory (LSTM) layers. The fully connected model part consisted of a linear layer, a leaky ReLU (rectified linear unit) activation function, a batch normalization layer, and a multi-sample dropout scheme^77^ followed by a sigmoid function.

We compared three different model architectures for the glycan analysis module. For assessing GlyCompare,^27^ the glycan analysis module comprised a fully connected neural network using the 12,259 GlyCompare features as inputs for two linear layers interspersed with dropout, leaky ReLU, and batch normalization layers. For the model containing a SweetTalk-based language model for glycan analysis,^22^ we converted glycans to glycowords and used a bidirectional recurrent neural network for protein sequences. For the SweetNet-based model,^24^ we converted glycans to graphs by constructing a list of nodes (representing monosaccharides or linkages) and edges to denote graph connectivity. All glycan processing for SweetTalk and SweetNet was done using glycowork version 0.5.^78^ The corresponding model contained an embedding layer and three graph convolutional layers, interspersed by leaky ReLUs, Top-K pooling layers, and both global mean and global maximum pooling operations. Model architectures and hyperparameters were optimized using cross-validation.

### Model training and prediction

All models were trained with an NVIDIA^®^ Tesla^®^ K80 GPU using PyTorch version 1.11.0.^79^ We split the data on a protein level into 80% for training and 20% for testing. For the RNNs, all glycosite-flanking protein sequence and glycan structure were brought to the same length by padding. Linear layers and RNNs were initialized using Xavier initialization^80^ while SweetNet-type models were initialized using a sparse initialization scheme with a sparsity of 10%.

We used a batch size of 64 for all models. As an optimizer, we used ADAM (adaptive moment estimation) with a weight decay value of 0.00001 and a starting learning rate of 0.00001, which was decayed according to a cosine function over 170 epochs. We trained models for a maximum of 250 epochs, with an early stopping criterion of 25 epochs without a decrease in validation loss. As a loss function, we used binary cross-entropy. Beginning from epoch 150, we additionally employed stochastic weight averaging^81^ with a learning rate of 0.0001.

The presence or absence of each glycan can be predicted from the trained InSaNNE model by inputting a glycosite and glycans to predict whether these glycans could occur on this glycosite. To heuristically boost signal for glycans with limited representation in the training set, we generated a naturalistic background of predicted presence for each glycan. Predictions were generated from all training-set glycosites to capture the biases and variation of the dataset as a background predicted-presence distribution for each glycan. The background-adjusted predicted-presence is the product of predicted presence and the predicted-presence cumulative probability (statsmodels::ECDF v0.12.2) relative to the naturalistic background for that glycan.

### Integration and display of predictions in GlyConnect

Using InSaNNe, we calculate the predicted presence of 512 N-linked glycans for each N-linked glycosite in the GlyConnect dataset. Prediction data were processed to fit the requirements of the GlyConnect database format, mainly storing association between glycans, glycoproteins and glycosites.^30^

IUPAC-represented glycans,^82,83^ output by InSaNNe, were transformed to GlycoCT^84^ using the GlyConnect API function, convertIupacToGlycoct (https://bitbucket.org/sib-pig/sugar-converter/downloads/). Transformed prediction data was integrated in the database to enable dynamic mapping through predefined queries for glycan structures and glycoprotein sites. Once transformed, any update of the InSaNNE prediction will easily be reflected in the database.

JSON files resulting from querying GlyConnect REST API are used for data export and display. A d3.js heatmap (https://d3-graph-gallery.com/heatmap) was selected as an appropriate data visualizer. The dimensions are defined as glycan structures/compositions and glycoprotein sites (designated by UniProt accession numbers and glycosylated amino acid sequence position). Heatmaps are created in three types of pages: (1) protein page featuring all glycan structures and compositions found attached to that protein, (2) structure page, featuring one structure and the many proteins on which they are found attached, and (3) composition page, featuring all matching glycan structures and the many proteins on which they are found attached. This data can be exported as csv files. Prediction data can also be visualized and compared using GlyConnect Compozitor.^44^

## Notes

### Summary of Updates

These revisions provide high resolution figures to correct the low resolution figures in the previous upload.

