## Supplemental Marterials for "Decoding glycosylation potential from protein structure across human glycoproteins with a multi-view recurrent neural network"

### Supplemental Figures

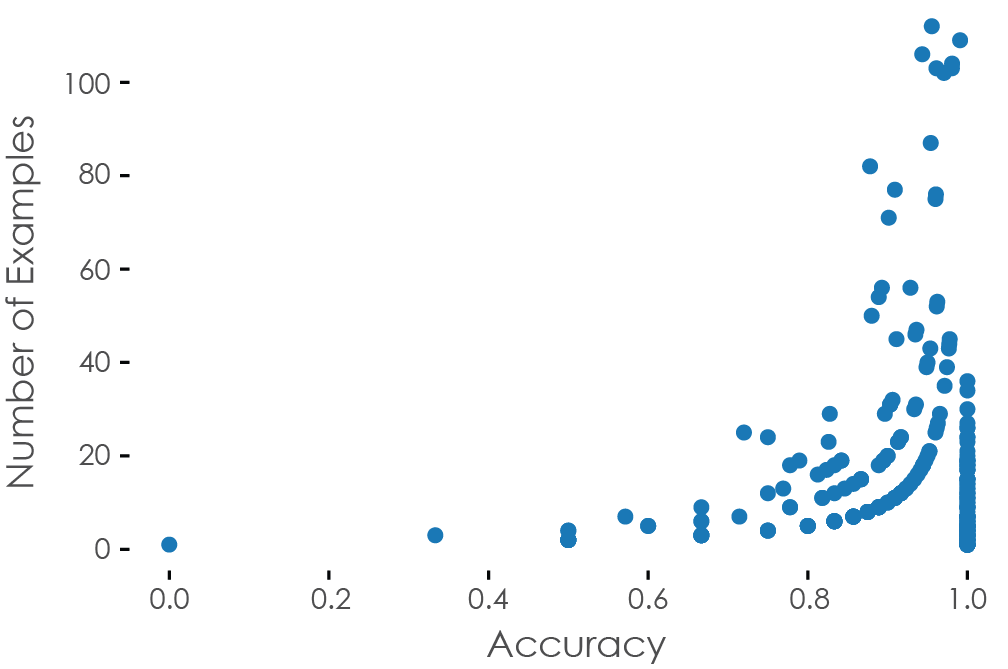

**Supplementary Figure** 1 **Dependence of sequon prediction performance on occurrence.** Using our trained InSaNNE model, we plotted the prediction performance of sequons, averaged over all their observed glycans, against the number of sequon-glycan pairs in our dataset.

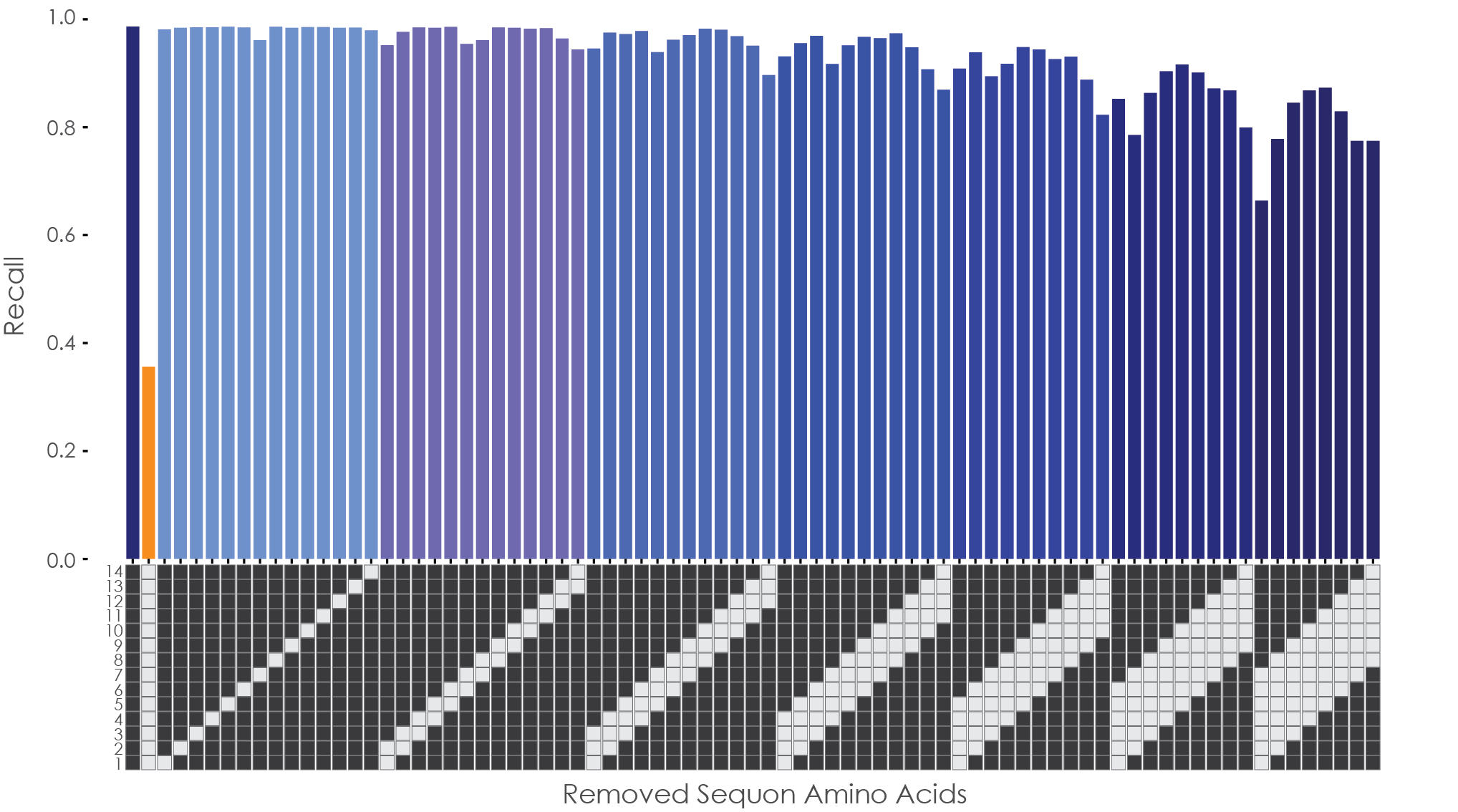

**Supplementary Figure** 2 **Redundancies in sequon sequence.** All sequon-glycan pairs in our dataset were used to obtain an averaged recall value for our trained InSaNNE model (WT), compared to an averaged recall value when removing the entire sequon sequence (all). We then iteratively removed single or multiple amino acid positions from all sequons and assessed model recall.

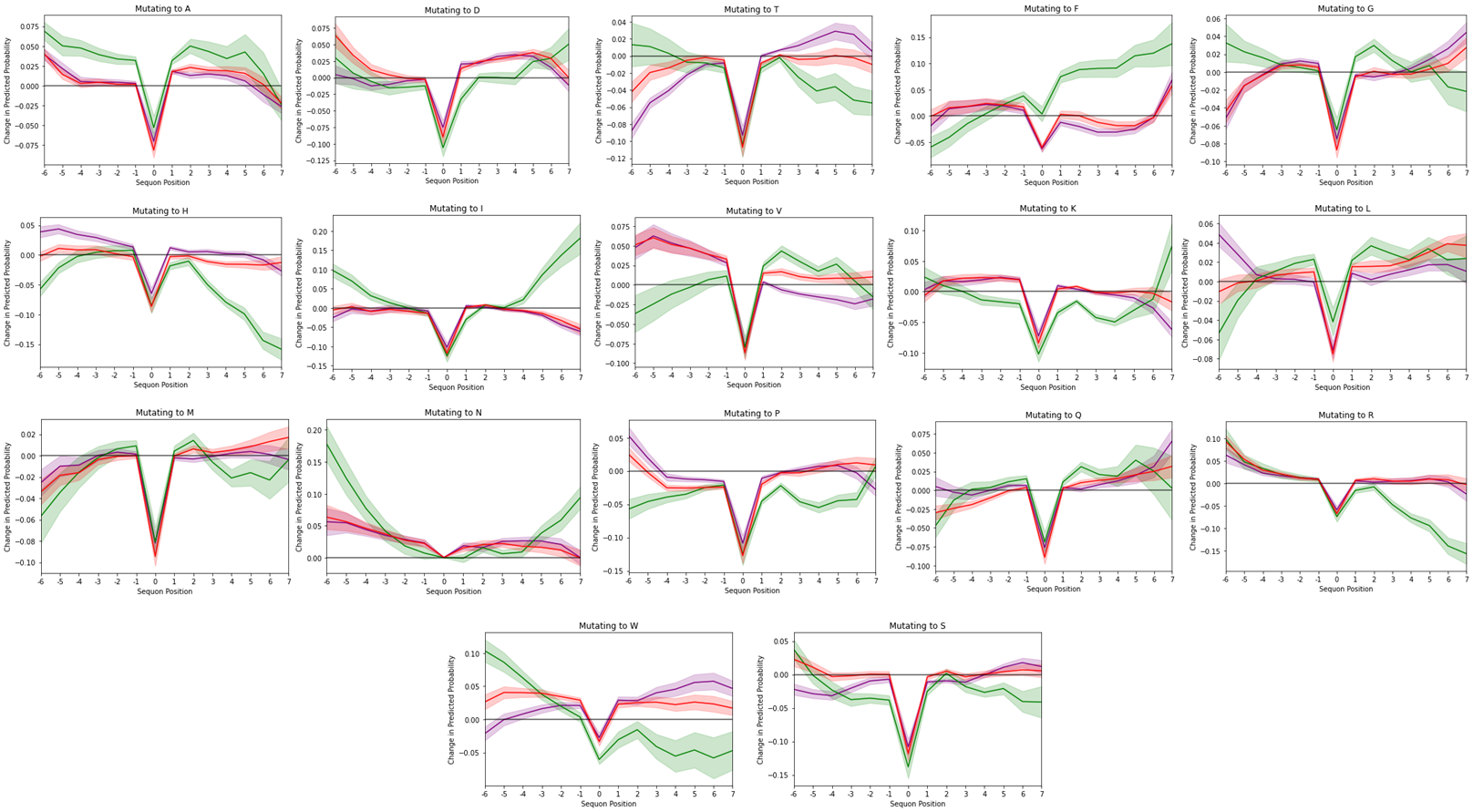

**Supplementary Figure** 3 **Effects of amino acid substitutions on predicted glycosylation ranges.** For all N-linked glycosites in our dataset, we systematically substituted all amino acids with every other amino acid and used the modified glycosite-flanking sequences as input for our trained InSaNNE model, obtaining a predicted glycosylation range. We then calculated the average change compared to the glycosylation range of the wild-type glycosites, which is shown here with the 95% confidence interval. Lines for changes to high-mannose (green), fucosylated (red), and sialylated (purple) glycans are shown, with a horizontal line at zero

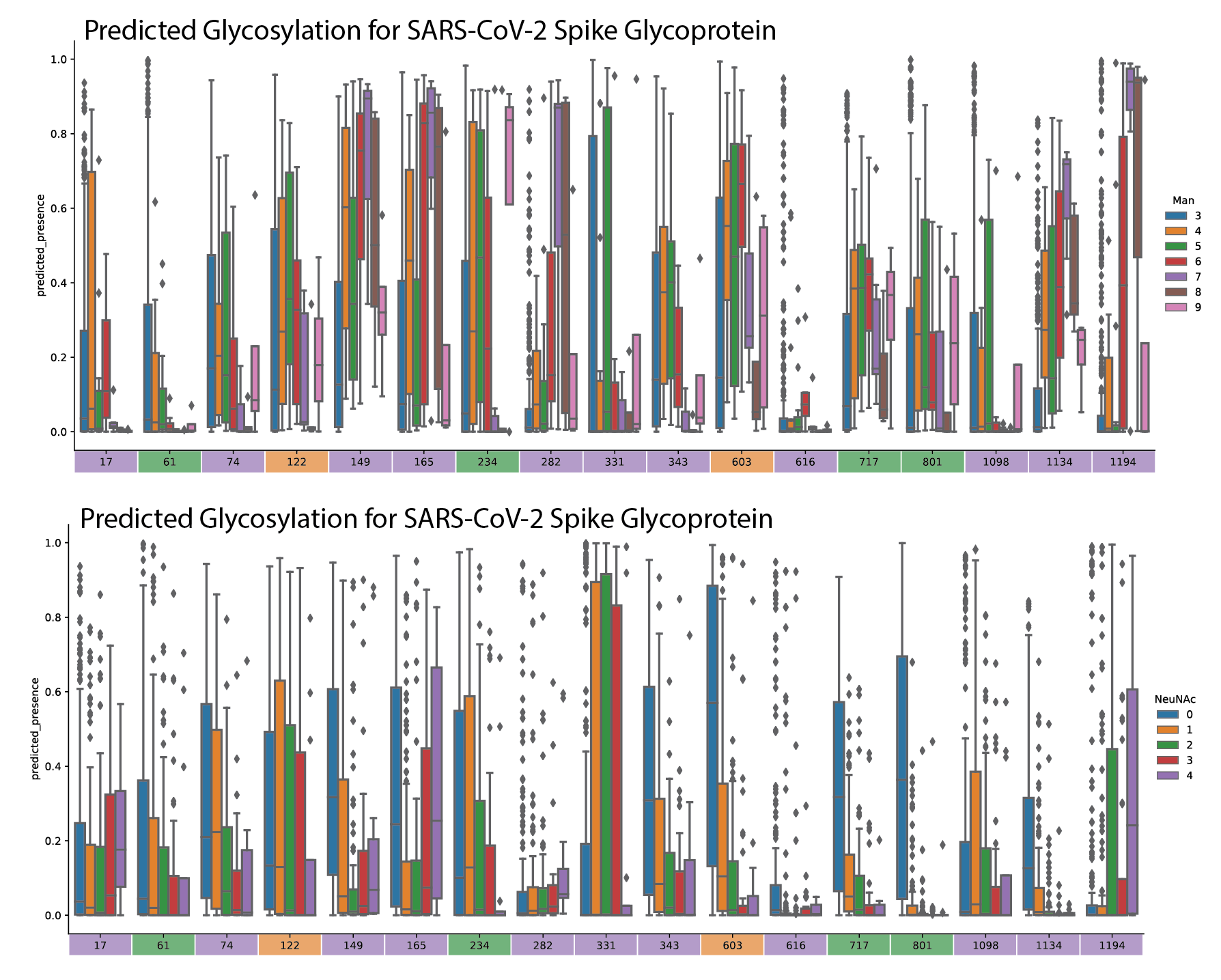

**Supplementary Figure** 4 **Predicted-presence by monosaccharide for all sites in the SARS-CoV-2 spike**. Glycosylation by mannose per glycan and sialic acid per glycan. Bottom colorbar in panels C and D indicate the dominant glycan type, complex (purple), hybrid (orange), oligomannose (green) characterized at that site in the wildtype spike ^15^.

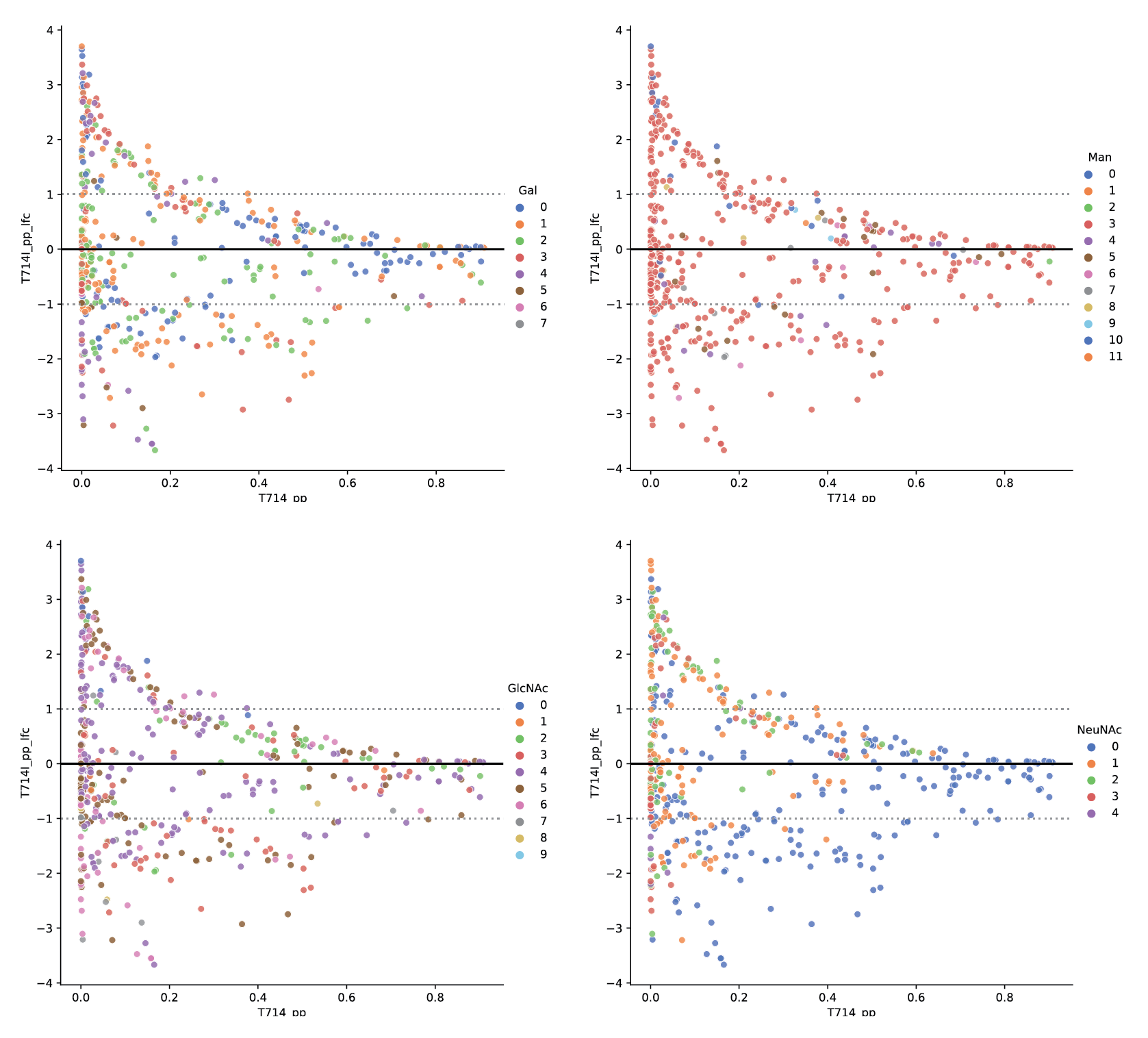

**Supplementary Figure** 5 **Predicted change in presence for glycans at N717 in B.1.1.7.** Predicted-presence fold-change (y-axis) is stratified by the basal predicted-presence for each glycan in the wild-type (x-axis). Predicted-presence fold-change from wild-type by galactose, mannose, GlcNAc, and sialic acid is provided for N717 B.1.1.7.

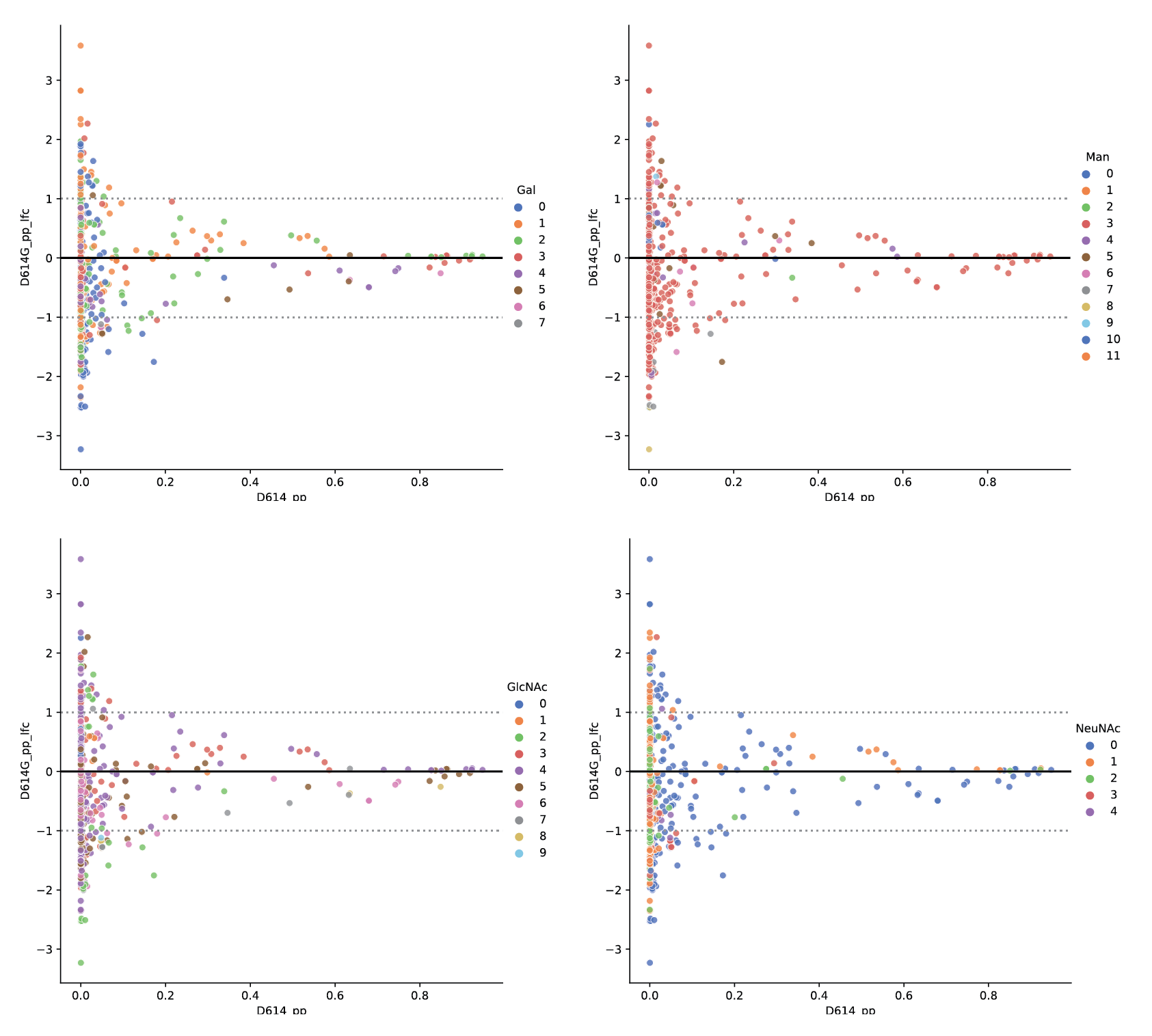

**Supplementary Figure** 6 **Predicted change in presence for glycans at N616 in D614G.** Predicted-presence fold-change (y-axis) is stratified by the basal predicted-presence for each glycan in the wild-type (x-axis). Predicted-presence fold-change from wild-type by galactose, mannose, GlcNAc, and sialic acid is provided for N616 in D614G.

### Supplemental Results

### Protein structure optimization and ablation demonstrates that all included feature types support predictive performance

To determine suitable dataset preparations, we trained a random forest classifier to distinguish oligomannose from complex glycosylation sites, given protein structure and surface data. Dataset preparations included protein structure model type (ab initio [I-TASSER], curated [SWISSMOD], or empirical [PDB]), and proximity radius defining “proximal” amino acids (4-10Å). Hyperparameters were optimized using 500 iterations of grid-search. Labels were balanced using up-sampling and performance was evaluated using area under the receiver-operator curve (AUROC), Sensitivity and Specificity on two iterations of six-fold cross-validation; each fold contained non-overlapping groups of proteins to avoid overfitting due to protein identity.

By varying these parameters, we evaluated the optimal protein model and annotation resolution (**Table 1**). Models trained on I-TASSER protein structures with a 6Å-annotation resolution showed the highest performance across all three metrics relative to random forest models trained on I-TASSER proteins annotated at other resolutions. Among models trained using PDB protein structures, those trained on data annotated at 8Å performed best across all three metrics. The best PDB-trained models measured, on average, comparable AUROC, a 3.5% decline in sensitivity and 13% decline in specificity compared to I-TASSER-trained models. SWISSMOD-trained models did not have a clear best resolution though average scores were mostly comparable to those trained using PDB or I-TASSER structures. Overall, the highest performing models appear to be those trained with either I-TASSER or PDB protein structures with resolution 6-8Å.

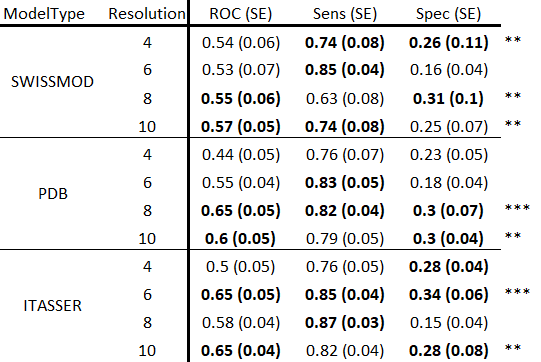

*Table 1 - Random forest model performance in a 2x6-fold cross-validation. Cross validation folds were split on protein identity. Training data were either annotated from SWISSMOD curated homology models, PDB empirical models, or ab initio I-TASSER homology models using structural resolutions between 4-10 Å. Performance was measured using average AUROC, Sensitivity and Specificity across cross-validation and the standard error across 12 folds in parenthesis. For each model type, the top two scores are bold. Rows with two or more top scores are noted two or more “*” in the final column.*

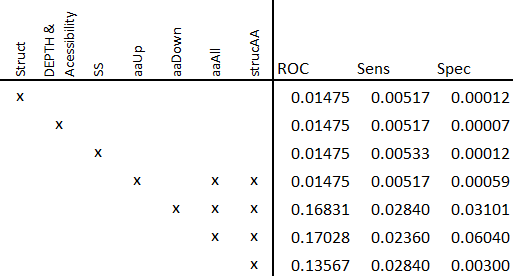

*Table 2 - Ablation of major protein structure feature types. Seven ablations of major feature-types including: struct (all structure-derived annotation), Depth & Accessibility (depth of residue, relative/absolute surface area), SS (secondary structure), aaUp/Down/All (sequence-proximal amino acids upstream/downstream/either), structAA (structure-proximal amino acids). Because some feature-types are associated, the ablation of some feature-types such as aaUp also required the removal of other feature-types indicated by the “x” left of the center line. Ablation significance is indicated by FDR-corrected Fisher’s Method pooled p-values (2-sample t-test, n=12 for each sample) comparing the performance distribution of ablation trained models to models trained on all data; performance distributions were collected over 2x6-fold cross-validation.*

Towards determining the importance of different protein structure annotations, we performed an ablation analysis by removing major types of data from the training set and comparing performance to models trained on all data (**Table 2**). We pooled the significance of each depletion in performance relative to models trained on all data (2-sample t-test, Fisher’s method for pooling p-values); p-values were pooled within each ablation and performance metric across models trained on I-TASSER, PDB and SWISSMOD protein structures at all resolutions. AUROC, sensitivity and specificity are all more sensitive to ablations in secondary structures, depth and upstream amino acids (FDR<0.01475). Interestingly, only sensitivity and specificity significantly decrease when downstream amino acid or structurally proximal amino acid data is removed (FDR<0.03). Overall, each major datatype is necessary to maintain performance across all three metrics.
